# NanoDel: Identification of large-scale mitochondrial DNA deletions using long-read sequencing

**DOI:** 10.1101/2025.09.19.677263

**Authors:** C. Fearn, J. Poulton, C. Fratter, C. Oliva, C. Griguer, R. A. Baldock, S.C. Robson, R.E. McGeehan

## Abstract

**Motivation:** Traditional methods for detecting large-scale mitochondrial DNA (mtDNA) deletions (LSMDs) in cells present challenges, i.e. *a priori* information, high DNA inputs, poor sensitivity and are not always quantitative. Mitigation can be achieved through high throughput DNA sequencing using e.g. Illumina and Oxford Nanopore Technologies (ONT), in combination with LSMD breakpoint identification and quantification using bioinformatic tools. Splice-aware RNA alignment tools increase the sensitivity for detecting LSMD breakpoints compared with DNA aligners. Long-read sequencing (LRS) also offers potential advantages over short read sequencing, e.g. greater read lengths and capturing variants on single reads. No existing pipelines capture the benefits of both a splice-aware alignment tool and LRS.

**Results:** We developed “NanoDel”, a LRS pipeline, to sensitively and accurately detect cellular LSMDs. Using artificial datasets, “NanoDel” was more sensitive and accurate than other pipelines. In samples diagnosed with mitochondrial disease, it identified both known and previously uncharacterised (including mixtures) of LSMDs, without *a priori* information. LSMD breakpoints were found in *mt-co1, mt-cyb, mt-nd6* and *mt-nd5* genes. Analysis of selected LSMDs revealed proximity to repeat and putative G-quadruplex motifs, and occurrence in a range of healthy and pathological tissues, indicating potential for a shared vulnerability landscape in mtDNA, shaped by sequence motifs and structural constraints. “NanoDel” combined with one-amplicon, not two-amplicon, LR-PCR offers a robust strategy with clinical application for detecting LSMDs across a variety of cell/tissue samples, and it’s application across a broader range of samples, will yield new mechanistic insights into LSMD formation, and further our understanding of mtDNA instability.

## Introduction

Mitochondria are cellular organelles that have their own double-stranded, supercoiled genome; mitochondrial DNA (mtDNA). MtDNA has 38 loci, including 13 protein coding genes, 2 rRNAs, 22 tRNAs and the control region (D-loop). The D-loop is integral for producing the multi-protein mitochondrial respiratory chain (MRC) complexes (I-IV), which along with complex V forms part of the oxidative phosphorylation (OXPHOS) system. This system, which also includes approximately 70 nuclear-encoded subunits, is central to ATP generation and cellular metabolism (reviewed in (Vafai and Mootha 2012)). Direct and/or indirect alterations in the mitochondrial genome can impact the function of the OXPHOS system, and therefore cellular energy generation, contributing to disease (for examples, see Table S1). Large-scale mtDNA deletions (LSMDs), which are direct alterations to mtDNA, result in genes crucial for cellular energy production via OXPHOS being removed and, therefore, primary mitochondrial disease.

Several traditional methods, such as Southern blotting (Moraes et al. 1989)(Santorelli et al. 1996), PCR (Schon et al. 1989) and Sanger sequencing (Johns et al. 1989)(Poulton, Deadman, and Gardiner 1989) (Johns et al. 1989) (Mita et al. 1990), have been used historically to detect LSMDs. However, the accurate breakpoint and heteroplasmy determinations using these techniques is challenging, requiring *a priori* breakpoint information, and high DNA inputs. Often such methods also lack sensitivity, are not completely quantitative, and can be labour intensive. One way to identify the breakpoints where multiple unknown LSMDs are present would be to plasmid clone and select all molecules prior to Sanger sequencing, however this is even more labour intensive (Schon et al. 1989).

A useful and sensitive alternative for identifying unknown LSMDs is by LR-PCR to amplify whole mtDNAs, followed by high-throughput DNA sequencing using platforms from e.g., Illumina (Hjelm et al. 2019) or Oxford Nanopore Technologies (ONT), and subsequent breakpoint identification and quantification using bioinformatic tools. Indeed, in 2019, Hjelm *et al*. published a pipeline that combines the LR-PCR of whole mtDNAs, as single amplicons, followed by Illumina sequencing, and a splice-aware RNA alignment tool for breakpoint identification and quantification. This demonstrated increased the sensitivity in detecting true positives compared to traditional DNA alignment tools (e.g. Burrows-Wheeler aligner) by around 60% (Hjelm et al. 2019). While examples of LMSDs detection by ONT sequencing exist, to date these have not used LR-PCR or a splice-aware aligner (Vandiver et al. 2023)(Frascarelli et al. 2023). However, employing LR-PCR and ONT sequencing for LSMD detection could offer additional advantages over short read (e.g., Illumina) sequencing, including less amplification of nuclear mitochondrial DNA segments (NUMTs) and longer individual read lengths (>1kb vs. <300bp) that span breakpoints, meaning a single read can represent an entire variant. This allows for the possibility of analysing variants, including SNVs, in tandem on the same read.

With improvements to sequencing chemistry and base-calling models now achieving quality scores above Q20, ONT is an attractive alternative to Illumina for the accurate breakpoint identification and quantification of LSMDs. However, the lack of an ONT pipeline comparable to that of (Hjelm et al. 2019), prompted us to develop “NanoDel”. This pipeline endeavours to amplify whole mtDNAs as either one or two amplicons by LR-PCR, followed by ONT sequencing, and a bespoke combination of bioinformatic tools for accurate breakpoint identification and quantification in cells/tissues.

In this study we first compared the ability of four pipelines to determine LSMDs breakpoints at different heteroplasmy levels using two artificial data sets. The four pipelines included two that use long-read sequences either in combination with a splice-aware RNA aligner (“NanoDel”, Pipeline 1, this study) or a DNA aligner (“Wood ONT”, Pipeline 3, (Wood et al. 2019)), and two that use short read Illumina sequences either in combination with a splice-aware RNA aligner (“Splice-Break”, Pipeline 2, (Hjelm et al. 2019)) or a DNA aligner (“Wood Illumina”, Pipeline 4, (Wood et al. 2019)). Second, we directly compared the two different ONT-pipelines (“NanoDel” and “Wood ONT”) to understand the impact of the selected alignment strategy, and the two LR-PCR approaches (one and two-amplicon) to determine known LSMDs in Kearns-Sayre syndrome (KSS) tissues. Third, we compared the LSMDs information generated by “NanoDel” and one-amplicon PCR against LSMDs information obtained diagnostically (using traditional methods) for additional patients with mitochondrial disease. This provides a comprehensive benchmarking of the power of NanoDel for exploring unknown LSMDs from clinical samples, allowing for fine-tuned understanding of the impact of mtDNA instability on disease.

## Methods and Materials

### Patient-derived tissues, cell cultures and DNA extraction

Three total DNA samples extracted from the kidney, cerebellum, and muscle of a KSS patient of known LSMD composition determined by Southern blotting (Table S2) were provided by Professor Joanna Poulton at the Nuffield Department of Women’s and Reproductive health (Patient 1 in (Poulton et al. 1993)(Poulton, Deadman, and Gardiner 1989)(Poulton et al. 1995)), University of Oxford. Additionally, DNA samples from muscle (muscle_RNASEH1, muscle_TWNK and muscle_TYMP) and cultured cells (cells_nd and fibroblasts_nd) were provided by Professor Joanna Poulton and by Carl Fratter at the Oxford Genetics Laboratories (OGL). These DNA samples were prepared and de-identified prior to transfer to our laboratory.

The original tissues were surplus to the quantity required for diagnosis, with the KSS patient tissues donated for research on the 1 January 1990 with permission from the patient’s family and surgeon, and the remaining patient tissue donated to research between 19 May 2005 and 13 December 2024 with written informed consent from all participants and with the approval of the United Kingdom National Research Ethics Service (South Central-Berkshire, #REC 20/04, IRAS 162181). Previous observations, i.e., LSMD composition determined by a range of different techniques following diagnosis at the OGL, were made available for these samples following sequencing analysis to prevent bias (Table S2). Details of cell lines used as mtDNA positive and negative amplification controls are provided in the Supplementary materials and methods. Total DNA was extracted from all cell culture samples using the QIAmp DNA mini kit as per the manufacturer’s instructions (Qiagen, 51304).

### Whole mitochondrial genome sequence generation

Whole mtDNAs were amplified from patient DNA in either two overlapping amplicons (Lloyd et al. 2012)(Lloyd et al. 2015)(Keatley et al. 2019) or as a single amplicon (Hjelm et al. 2019) by LR-PCR using human mtDNA-specific primers from total DNA (Table S3). Briefly, Nanopore libraries were prepared from either two or single LR-PCR amplicons using the Native Barcoding Kit 96 V12 library kit (ONT, NBD112.96), barcoded using the NBD112.96 kit and the NEB Blunt T/A Ligase MasterMix (NEB, M0367S), multiplexed and then cleaned using AMPure XP beads (Beckman Coulter, A63880). Adapter sequences were ligated to the multiplexed DNA sample using the NEBNext Quick Ligation module (New England Bioscience, E6056S). The prepared, washed and quantified DNA library was then loaded to an R10.4 flow cell and sequenced for 72 hours on an Oxford Nanopore Technologies GridION X5. Full details of the one- and two-amplicon LR-PCR, library preparation and sequencing are provided in the Supplementary materials and methods.

### Bioinformatics, artificial data generation, alignment and LSMD analyses

First and briefly, two styles of artificial FASTQ format raw sequence data files were generated using VISOR (v1.1, (Bolognini et al. 2020) in: 1) an ONT style with a longer average read length of ∼8kb and a “Nanopore 2020” style error model, and 2) an Illumina style with 250bp length reads and an “Illumina” style error model. Each style contained either controlled quantities of reads designed to emulate a single deletion (mtDNA 4977) at different levels; i.e., 75%, 50%, 25%, 5%, 1% and 0.25% of reads (artificial data 1) or multiple deletions (m.8471-13449del, m.6620_8440del, m.3451_5500del and m.12538_14572del) at a fixed quantity (10%) of reads (artificial data 2). These files were then used to evaluate and compare the efficacy of the four separate alignment and deletion calling strategies outlined above (Pipelines 1-4). Second, raw reads from the patient mtDNA LR-PCR libraries (one- and/or two-amplicon) were base-called and demultiplexed (including barcode trimming) using Guppy (v6.2.7 and v6.3.9) and transferred to a local high-performance computing cluster running on Ubuntu (v20.04.5). Base-called reads were initially filtered to remove reads of <20bp in length or a Q-score <10 using NanoFilt (v2.8.0, (De Coster et al. 2018) prior to processing through the two best performing pipelines: “NanoDel” (Pipeline 1) and “Wood ONT” (Pipeline 3). In this study all reads (artificial and patient) were aligned to the rCRS reference sequence (NC_012920.1, (Bolognini et al. 2020)(Andrews et al. 1999)). Full details of the artificial data generation, alignment and deletion calling strategies, including all read quality control, software, benchmarking and filtering are given in the Supplementary materials and methods.

### Repeat and G4 sequence, previous tissue/ disease association and statistical analyses

Sequences flanking the 5’- and 3’-breakpoints of the mitochondrial disease patient LSMDs occurring at ≥50% of reads detected in the filtered “NanoDel” data (Pipeline 1) were analysed for repeat sequences and their potential to form G-quadruplexes (G4s) using QGRS mapper (Kikin, D’Antonio, and Bagga 2006)) and AlphaFold 3 (AF3)(Abramson et al. 2024)).

Previous tissue and/or disease associations of the LSMDs, both “NanoDel” (Pipeline 1) and adjusted positions (Table S6), and 10bp flanking either side were investigated using the Mitobreak database, which contains 1369 human mtDNA deletions previously described in the literature (Damas et al. 2014).

Statistical analyses of post-alignment read metrics were conducted using GraphPad Prism (v9.4.1). The normality of each metric for each pipeline was assessed using the Shapiro-Wilk test (Shapiro and Wilk 1965). Where assumptions of normality were met, a Brown-Forsythe and Welch ANOVA was employed to model differences between the approaches. If the assumption of normality was violated, the non-parametric Kruskal-Wallis test (Kruskal and Wallis 1952) was used instead. Further post-hoc pairwise comparisons were made using either Welch’s t-test or the non-parametric Mann-Whitney test, respectively. Full details on the repeat and G4 analyses are given in the Supplementary materials and methods.

## Results

The pipeline developed for this study, “NanoDel” (Pipeline 1), was tested against three other pipelines using two artificial datasets to compare the performance in detecting the percentage of reads within which each LSMD occurs, and its location. NanoDel offered the best balance between accuracy, sensitivity and specificity when identifying the heteroplasmy and breakpoint location, both as a single deletion (mtDNA4977) in artificial data set 1, and as mixed deletions in artificial data set 2 (Fig 2-3, and Supplementary Results).

**Fig 1.**
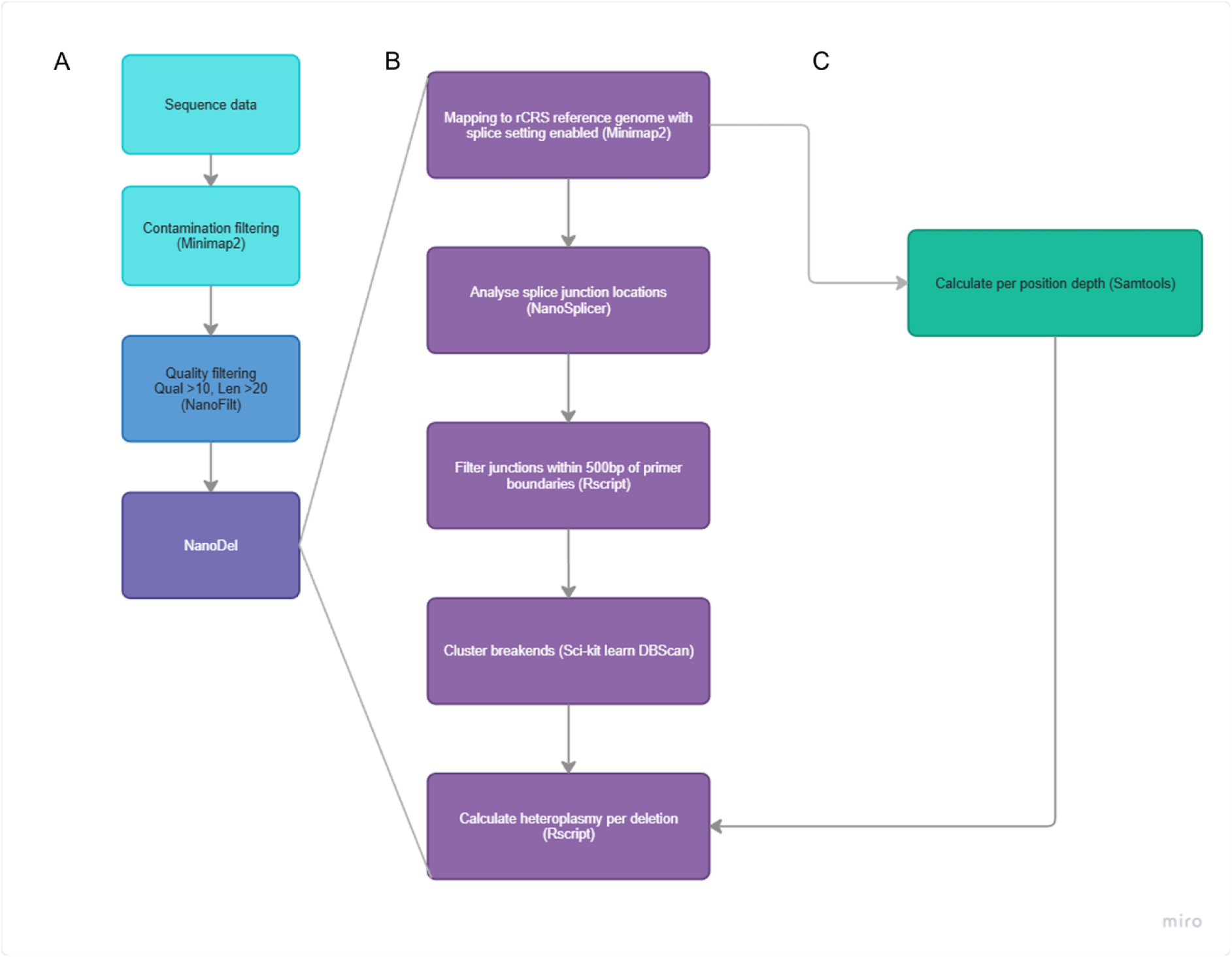
Flow diagram of large-scale mtDNA deletions using Pipeline 1 (“NanoDel”) developed for this study. A) Raw read processing. Raw reads were base-called following barcode trimming and demultiplexing at run-time in MinKnow using Guppy (v6.2.7 or v6.3.9) and transferred from the GridION to a local high-performance computing cluster (HPC) running on Ubuntu (v20.4.5). Where the version 14 chemistry was used, samples were aligned to the bacteriophage lambda reference genome (NC_001416.1) using minimap2 (v2.21) to remove reads mapping to the spike-in DNA control. All mapped reads were then subsequently removed from their constituent FASTQ files. Individual FASTQ files were then filtered to remove all reads <20bp and/or with a Q-score <10 using NanoFilt (v2.8) post-barcode trimming, but before being processed through “NanoDel”. B) “NanoDel” was constructed and run on the HPC cluster with Ubuntu (v20.4.5). First, filtered reads were aligned to the revised Cambridge reference sequence (rCRS, NC_012920.1, Andrews et al., 1999), using Minimap2 (v2.21), with splice alignment settings enabled (-ax splice), but preferences for canonical splice junctions disabled (-u no). These outputs are piped directly to SAMtools, where the resulting data were converted to BAM format using SAMtools view (-S and -b options for SAM format input and BAM output) and sorted using the SAMtools sort function. Splice junction locations were then extracted from the BAM files using the JWR_Checker.py script from NanoSplicer (v1.0), with output CSV format options enabled (–output_csv). Simultaneously, the per-base depth was calculated by the SAMtools depth function for all positions regardless of coverage (option -aa) and later used for heteroplasmy calculation. Custom R scripts were then available as options within the pipeline (filt or no_filt) that either reformatted the data and removed any deletion junction calls with 5’- or 3’-breakpoints within 500bp of a primer binding site, or not. This is because (Hjelm et al. 2019) demonstrated an increase in false positive breakpoint calls within this distance of primer boundaries. The remaining breakpoint junctions were then clustered using a custom python script that employed the sci-kit learn package (v1.2). A density-based clustering approach (DBScan) was used to group all LSMD calls with both 5’- and 3’-breakpoints that were within 10bp of other 5’- and 3’-breakpoints. Average 5’- and 3’-locations were then taken from each cluster and reported with the number of reads generating that cluster for each sample and output in CSV format, with all points not contributing to a cluster being subset as ‘noise points’. C) Finally, a mean coverage benchmark was calculated from the depth output between positions m.500_1000 (according to the rCRS reference sequence) and used in the calculation of the relative read % (a proxy for heteroplasmy) for each LSMD using a custom R-script. As such the number of reads containing each specific LSMD was expressed as a percentage of the coverage benchmark for each respective sample.

**Fig 2.**
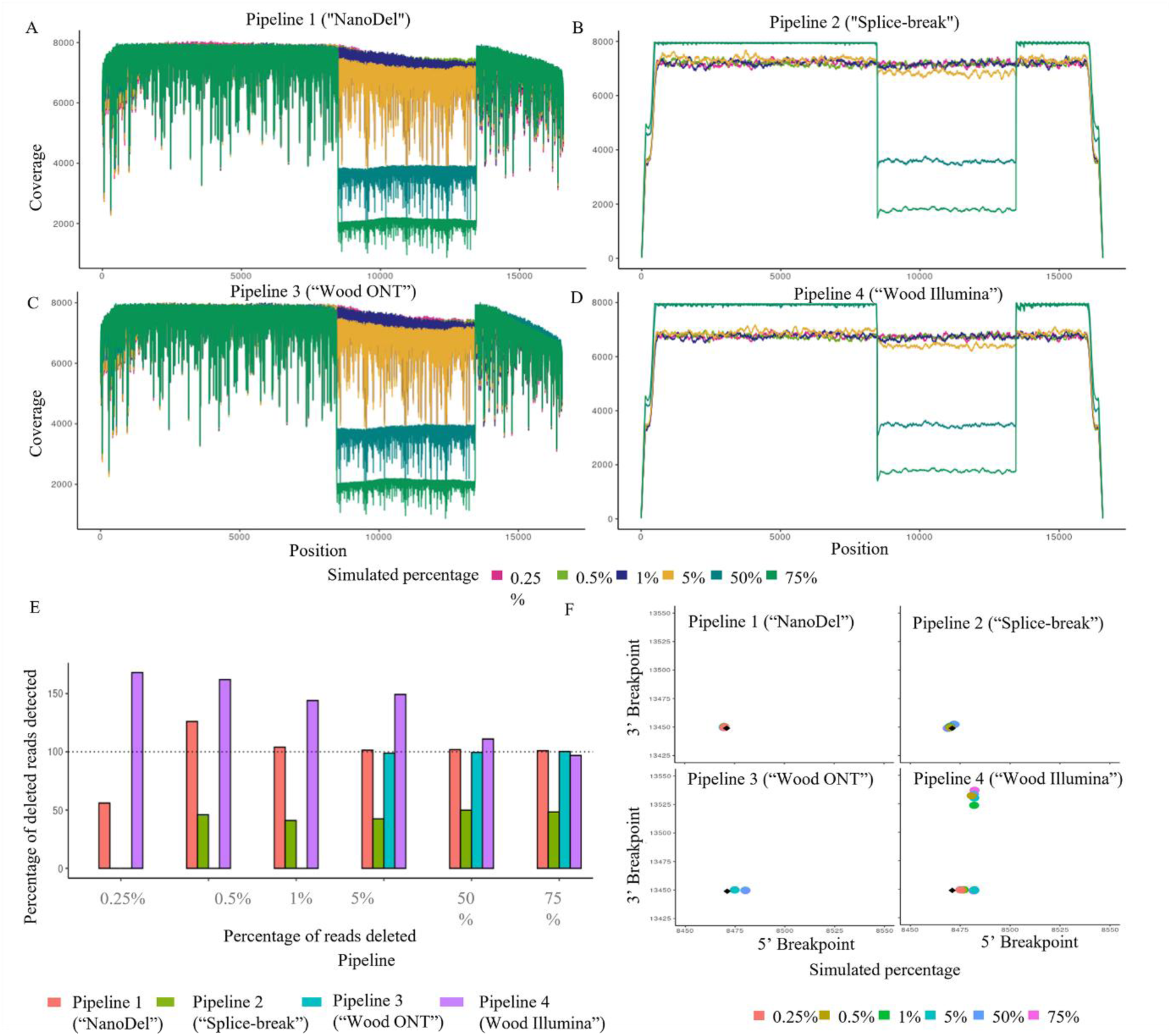
Analysis of artificial data set 1 that contains a mixture of the mtDNA4977 (m. 8471_13449del) and wild type mtDNA at different ratios. A-D) The per-position levels of coverage generated by Pipeline 1 (“NanoDel”), Pipeline 2 (“Splice-break”), Pipeline 3 (“Wood ONT”), and Pipeline 4 (“Wood Illumina”), respectively. E) The percentage of the expected number (7500 (75%), 5000 (50%), 500 (5%), 100 (1%), 50 (0.5%) and 25 (0.25%)) of deleted reads detected out of 10,000 identified by each pipeline. F) Breakpoint mapping locations identified at each deleted read% level (75%-0.25%) in relation to the true breakpoint locations (highlighted in black).

**Fig 3.**
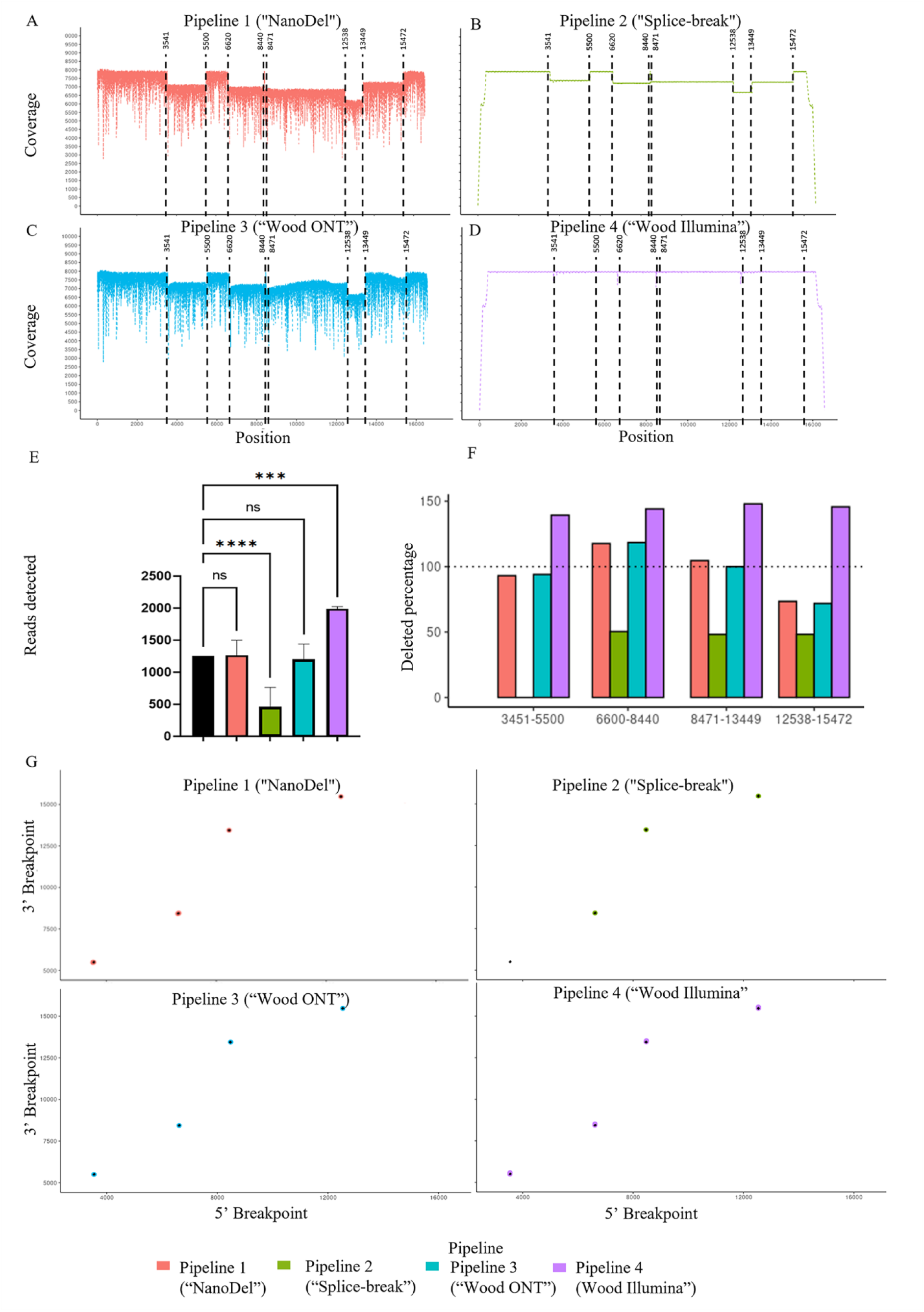
Analysis of artificial data set 2 that contains a mixture of deletions (i.e., between positions m. 8471_13449del, m. 6620_8440del, m. 3451_5500del and m. 2538_14572del), set at a read level of 10% and 90% wild type mtDNA. A-D) The per position levels of coverage generated by Pipeline 1(“NanoDel”), Pipeline 2 (“Splice-break”), Pipeline 3 (“Wood ONT”), and Pipeline 4 (“Wood Illumina”), respectively. Breakpoint locations are denoted by black dashed lines. E) The average number of deleted reads detected by each pipeline; each one is simulated out of 1250 reads. The percentage deleted reads detected by deletion reads (so reads detected out of 1250*100). Breakpoint mapping locations of each deletion identified in relation to the true breakpoint locations (highlighted in black). * = P<0.05, ** = P<0.01, *** = P<0.001, and **** = P<0.0001.

Due to the performance of “NanoDel” (Pipeline 1) in detecting LSMDs within the artificial data, it was then employed to detect LSMDs and determine their breakpoints in two small mitochondrial disease patient cohorts (Table S2) following both one- and two-amplicon LR-PCR. MtDNA breakpoints were also determined following LR-PCR amplification in these samples using “Wood ONT” (Pipeline 3) for comparison. LSMDs were observed in clinically diagnosed mitochondrial disease samples on agarose gels following different LR-PCR approaches, revealing single or multiple deletion types (Figure S1, and Supplementary results).

Post-alignment, “NanoDel” yielded lower error rates, higher mean coverage and higher maximum read lengths compared to “Wood ONT” (Pipeline 3). Furthermore, “NanoDel” (Pipeline 1) and “Wood ONT” (Pipeline 3) had increased mapping rate following one- rather than two-amplicon LR-PCR (Fig 4, Table S4 A-B, and Supplementary results).

**Fig 4.**
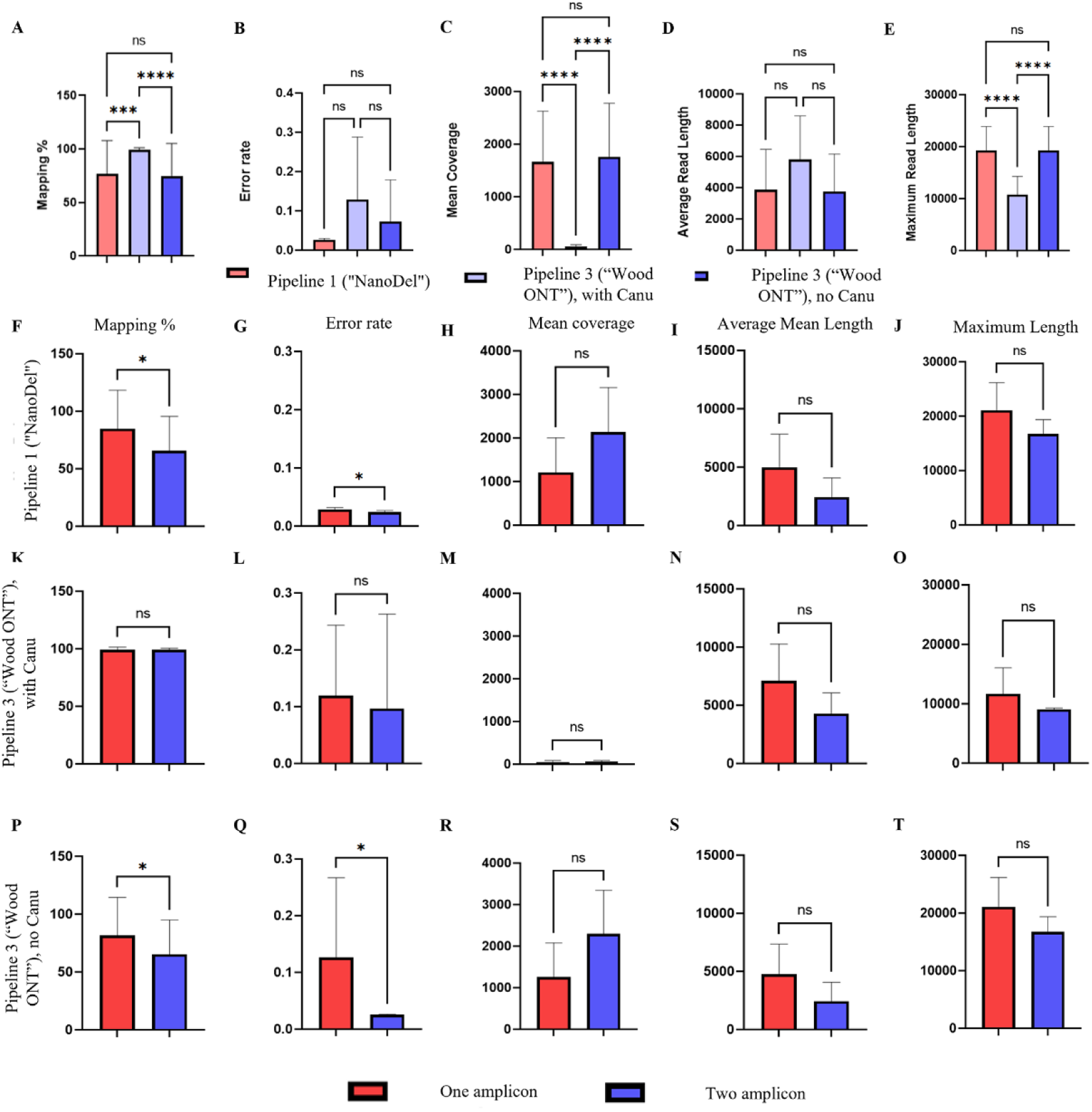
A comparison of post-alignment quality metrics: A, F, K and P) The percentage of reads mapped to the reference genome, B, G, L and Q) Error rate of mapped reads, C, H, M and R) The average mean coverage, D, I, N and S) Average read length, and E, J, O and T) Maximum read length following the alignment of one and two-amplicon LR-PCR reads, analysed in combination (A-E) or separately (F-T), by Pipeline 1 (“NanoDel”), Pipeline 3 (“Wood ONT”, with Canu) and Pipeline 3 (without Canu) generated from patients with clinically diagnosed mitochondrial diseases. * = P<0.05, ** = P<0.01, *** = P<0.001, and **** = P<0.0001.

In addition to the differences between the post-alignment quality metrics, “NanoDel” (Pipeline 1) and “Wood ONT” (Pipeline 3) in combination with one-amplicon, rather than two-amplicon, LR-PCR best validated a known LSMD (m.6130_15055del) in KSS patient tissues (Fig 5, Table S5A, and Supplementary results). In the two-amplicon LR-PCR data, deletion calling was impacted across amplicon boundaries, resulting in an impact to the LSMDs called from both the quality of the DNA and filtering applied to the variant calling.

**Fig 5.**
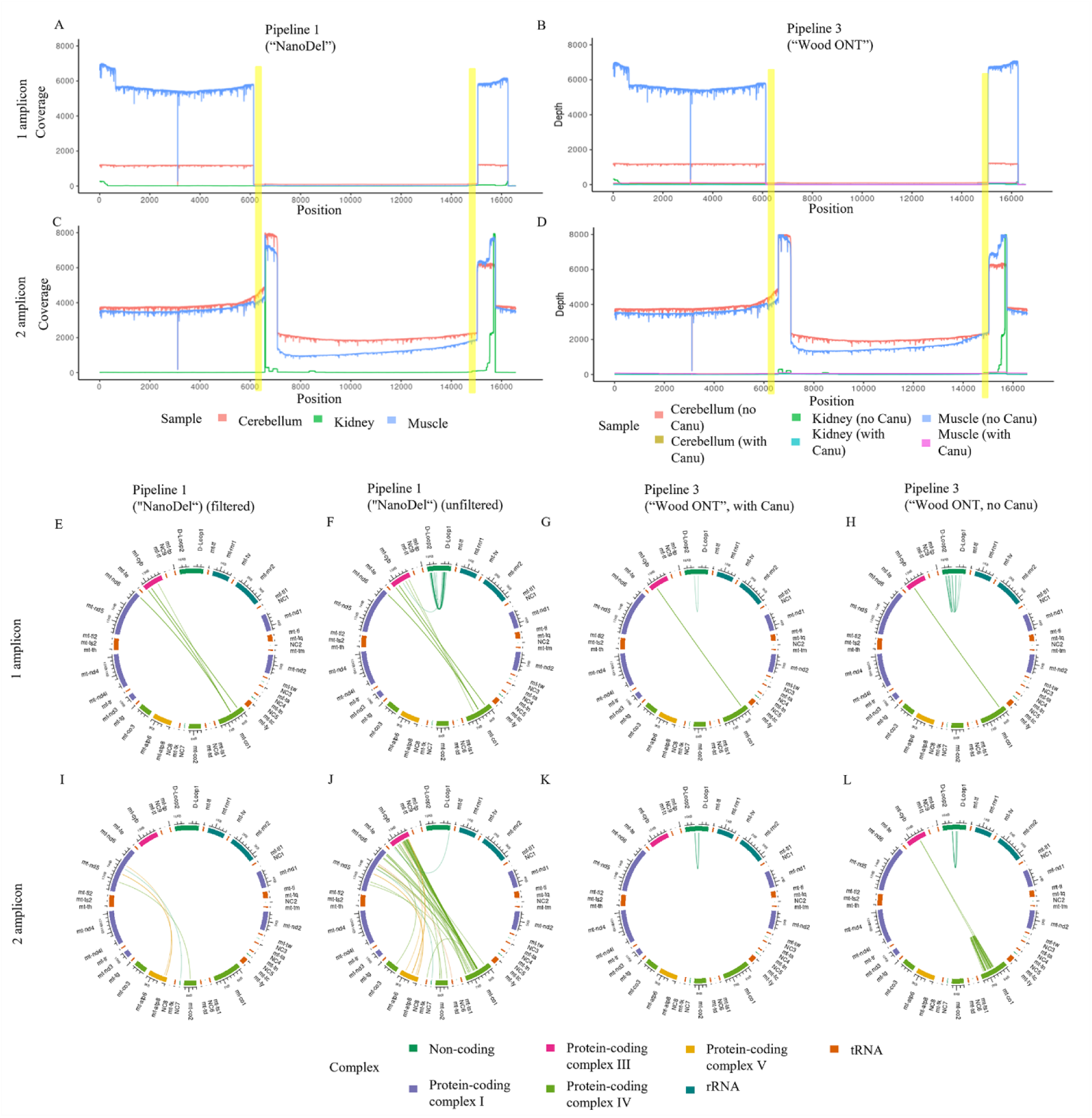
Analysis of coverage and deletion boundaries in different tissues of a KSS patient. A-D) The levels of coverage displayed as read depth at each position for each sample amplified as either one A, B) or two C, D) amplicons, aligned with Pipeline 1 (“NanoDel”, A, C) or Pipeline 3 (“Wood ONT”, B, D). E-L). Circos plots showing the 5’- and 3’-breakpoints of deletions following one E-H) and two I-L) amplicon LR-PCR aligned with Pipeline 1 (“NanoDel”) post-filtering E, I) and unfiltered F, J) and Pipeline 3 (“Wood ONT”) with G, K) and without H, L) Canu. The genomic links highlight the connected 5’- and 3’-breakpoints coloured by sample of detection, and key complexes across the mitochondrial genome are highlighted. Links proceed in a clockwise direction starting from the D-loop 1 region. Putative G4-structure forming regions are highlighted in yellow.

As the two-amplicon LR-PCR completely failed to detect the known deletion (m.6130_15056del) in the KSS samples (and could not detect deletions across amplicon boundaries), subsequent analyses of the other mitochondrial disease samples focussed on “NanoDel” (Pipeline 1) and “Wood ONT” (Pipeline 3) using just the one-amplicon LR-PCR. “NanoDel” (Pipeline 1) and one-amplicon LR-PCR validated known information obtained using traditional methods but also yielded new insights into LSMDs in the other mitochondrial disease patients, consistent with, or even in the absence of, primary diagnosis (Fig 6, Table S5B, and Supplementary results).

**Fig 6.**
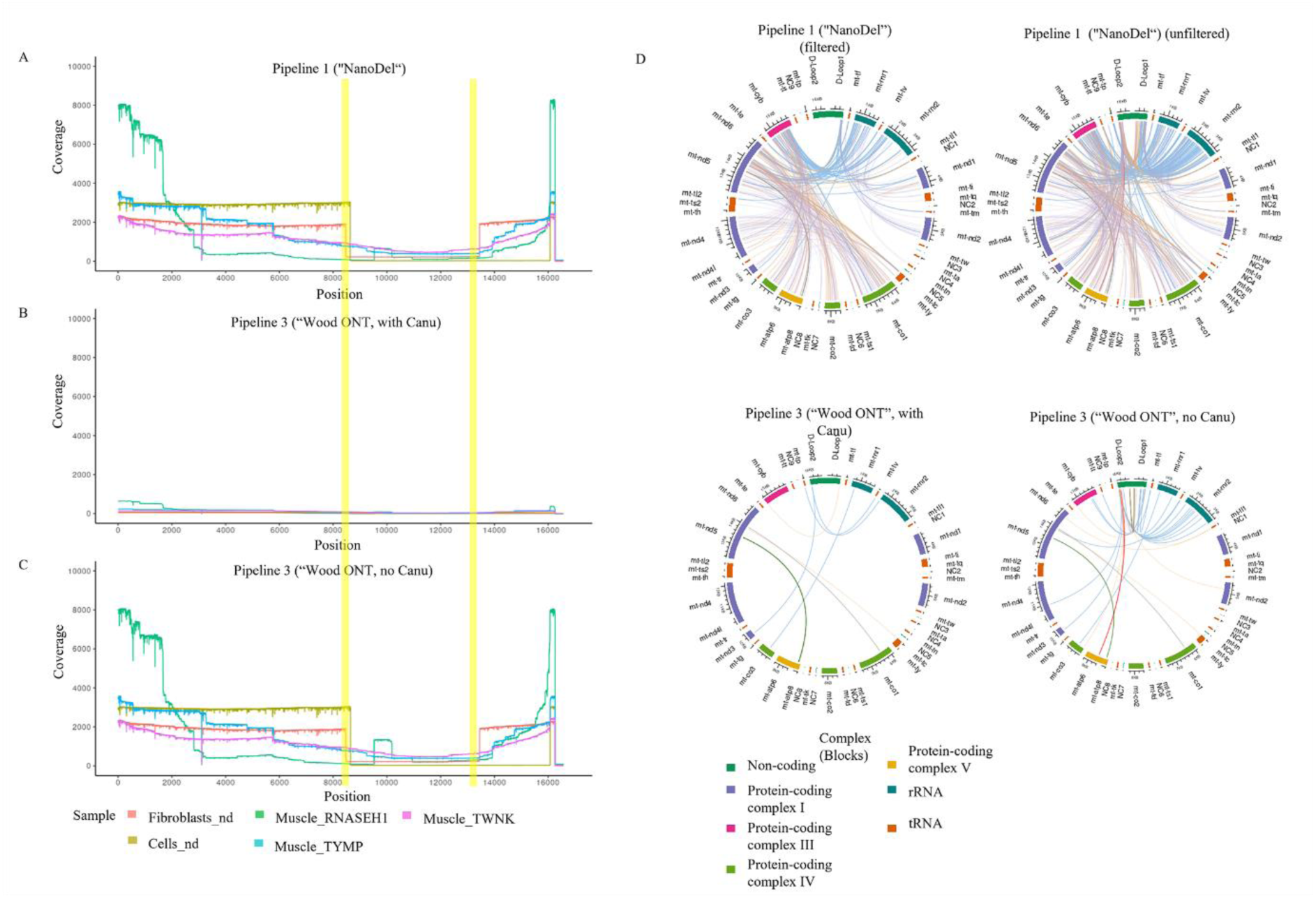
Analysis of coverage and deletion breakpoints in five samples clinically diagnosed with mitochondrial disease. A-C) Shows the coverage at each position for each sample generated with Pipeline (“NanoDel”, A), and Pipeline 3 (“Wood ONT“) with Canu, B) and without Canu C), respectively. D-G) Shows the 5’- and 3’-breakpoints of deletions identified by each pipeline in each sample as a circos plot following one-amplicon LR-PCR aligned with Pipeline 1 (”NanoDel“) post-filtering D) and unfiltered E) and Pipeline 3 (“Wood ONT”) with F) and without G) Canu. The genomic links highlight the connected 5’ and 3’ breakpoints coloured by sample of detection, and key complexes across the mitochondrial genome are highlighted. Links proceed in a clockwise direction starting from the D-loop 1 region. Putative G4-structure forming regions are highlighted in yellow.

Specifically, “NanoDel” (Pipeline 1) in combination with one-amplicon LR-PCR determined that *mt-co1, mt-nd5, mt-nd6* and *mt-cyb* harbour breakpoints most frequently in mitochondrial disease samples, and that breakpoints identified for m.6130_15055del in the KSS samples and m.8469_13446del in fibroblast_nd, which occurred at ≥50% of reads in filtered “NanoDel” (Pipeline 1) data lie adjacent to or within repeat sequences and putative G4 structure-forming sequences, and are associated with a range of healthy and pathological tissues (Fig 6, Table S6, and Supplementary results).

## Discussion

### Development, validation, and potential clinical application of NanoDel

In this study, we first developed and then evaluated the ability of four pipelines to accurately detect the location and quantify the read% of LSMDs using artificial data sets. These pipelines included both DNA and splice-aware RNA alignment methodologies, and were designed based on either short read (Illumina) or long-read (ONT) data sets. The pipelines selected for comparison against “NanoDel” (Pipeline 1, ONT splice-aware alignment, this study) were: “Splice-Break” (Pipeline 2, Illumina splice-aware alignment, (Hjelm et al. 2019)), “Wood ONT” (Pipeline 3, ONT DNA alignment, (Wood et al. 2019)) and “Wood Illumina” (Pipeline 4, Illumina DNA alignment, (Wood et al. 2019)). Splice-Break was chosen as a comparator as it uses Illumina short read sequencing in combination with a non-traditional splice-aware RNA alignment strategy that is not negatively affected by the presence of large-scale deletions, shown previously to improve LSMDs calling accuracy (Hjelm et al. 2019). “Wood ONT” (Pipeline 3) and “Wood Illumina” (Pipeline 4) were chosen as they have previously been tested in a UK diagnostic setting and use either Illumina short- or ONT long-read sequencing, respectively, in combination with a more traditional DNA alignment strategy (Wood et al. 2019).

We demonstrate that our pipeline, “NanoDel” (Pipeline 1), which calls LSMDs from long-read data using a splice-aware alignment approach, offers improvements in accuracy and sensitivity over the other publicly available tools. This is true in terms of both breakpoint location and read% quantification of mtDNA4977 (m.8471_13449del, (Yusoff et al. 2020)) and mixed populations of deletions, including at low levels, in artificial datasets. “NanoDel” (Pipeline 1) identified the correct breakpoint locations and number of reads for all the LSMDs. “Wood ONT” (Pipeline 3, DNA alignment) performed most similarly to “NanoDel” (Pipeline 1). This may be in part due to two factors: their use of a shared alignment tool ‘Minimap2‘ (albeit using different settings, with “NanoDel” making use of the splice preset settings); and the fact that they both analysed the same style of long-read data compared to the other pipelines analysing ‘short-read style’ simulated data. In comparison, both “Splice-break” (Pipeline 2, Illumina RNA alignment) and “Wood Illumina” (Pipeline 4, DNA alignment) could not identify the LSMDs at the lowest 0.25 read% levels (“Splice-break”), or identified false positives (”Wood Illumina”), and were the least able to accurately quantify the correct number of reads containing LSMDs. Both these pipelines made use of short Illumina-style reads.

While these findings align with separate prior reports emphasizing the importance of using splice-aware aligners in LSMDs calling (Hjelm et al. 2019) and the potential utility of ONT sequencing in mitochondrial genomics (Wood et al. 2019) (Frascarelli et al. 2023)(Vandiver et al. 2023) (Frascarelli et al. 2023), they also highlight the potential strength of combining splice-aware alignment methodologies with long-read sequencing (i.e. “NanoDel”) in LSMDs calling. Consequently, we wanted to deploy “NanoDel” (Pipeline 1), and “Wood ONT” (Pipeline 3) as a comparator, to investigate LSMDs in clinical samples from patients diagnosed with mitochondrial disease and varying amounts of LSMDs and compare with the information obtained using traditional techniques such as Southern blotting.

In doing so, we found that “NanoDel”, in combination with one rather than two-amplicon LR-PCR, outperformed the alternative “Wood ONT” in terms of mapping and error rate when investigating LSMDs in different tissues from mitochondrial disease patients. “NanoDel” in combination with one-amplicon LR-PCR was able to identify the previously observed LSMD (m.6130_15056del) (Poulton et al. 1995)(Poulton et al. 1993) (Poulton, Deadman, and Gardiner 1989) in KSS patient samples in a tissue specific manner. Furthermore, “NanoDel” was not only able to identify previously observed LSMDs in different tissues from five other patients diagnosed with mitochondrial disease, but also revealed additional, previously unreported LSMDs information. This demonstrated that “NanoDel” has power to identify LSMDs in different sample types (tissues/diseases), including more complex biological samples (such as those without a known genetic cause), with significant potential in a clinical setting.

Our findings demonstrated that “NanoDel”, in combination with one-amplicon LR-PCR, outperformed our two-amplicon approach. This is consistent with an original report that detailed the development and potential benefits of the one-amplicon LRPCR approach over multi-amplicon approaches, including increased discriminating power against nuclear mitochondrial DNA segments (NUMTs), more even coverage, and increased LSMD detection resolution by limiting amplicon dropout (Zhang, Cui, and Wong 2012). NUMTs are pieces of mtDNA that have been transferred to and integrated into the nuclear genome and can cause false positives during PCR amplification. Consistent with the original report, we also showed that NUMTs with “NanoDel” were not amplified in our rho-zero mtDNA depleted controls with either one- or two-amplicon LR-PCR. Further, we showed the coverage was more even in our samples, with one rather than two-amplicon LR-PCR. We found that LSMDs were only identified within individual amplicon boundaries with two- rather than one-amplicon PCR. Furthermore, the inflated number of reads within overlapping regions relative to benchmark coverage affected the read% of LSMDs. These findings suggest prior reports detailing mtDNA variants following the use of multi-amplicon PCR approaches, of which there are numerous examples including (but not restricted to) colorectal cancer (Aikhionbare et al. 2004), pancreatic cancer (Lam et al. 2012) and hypertension (Elango et al. 2011), could be incomplete and contain inaccuracies and should be interpreted with caution.

While the clinical application of ONT sequencing for mtDNA variants (including LSMD detection) has been proposed previously (Wood et al. 2019), recent improvements to the ONT chemistry now routinely generate base-calling quality scores of Q20+. This, taken together with “NanoDel”’s ability to yield a sensitive and accurate, and thus a comprehensive view of the LSMDs breakpoints/landscape without *a priori* information in individual samples (in combination with one-amplicon LR-PCR), offers improvements over traditional methods in this context. However, the limited of correlation in read% between “NanoDel” (and “Wood ONT”), in combination with one-amplicon LR-PCR, and the heteroplasmy measures obtained using traditional methods, could be due to the preferential amplification of shorter fragments during PCR in our samples (Kopsidas et al. 2000). While such preferential amplification has been shown to be minimised via the use of LR-PCR (Kopsidas et al. 2000), its impact could be verified in our samples by determining the ratio of deleted molecules to wild type molecules using digital PCR. Thus, for now, the read percentages should not be interpreted as heteroplasmy rates directly. Alternative PCR-free methods, e.g., purifying mitochondria/mtDNA prior to sequencing or WGS in combination with adaptive sequencing (where ONT pores screen for ‘expected’ sequences *in silico*, and reject strands that do not meet the expected requirements) may be used to increase sequencing yield of an intended target like mtDNA (Martin et al. 2022), and may more accurately quantify heteroplasmy. However, these methods also present challenges (e.g. Herbert et al. 2025). It can be difficult to obtain suitable yields of purified mtDNA (often due to its fragility), and WGS can generate poor coverage as the probes are not explicitly designed for mtDNA.

### Mechanistic insights and broader implications

Using “NanoDel”, we identified increased numbers of 5’-breakpoints in the gene *mt-co1* and 3’-breakpoints in the genes *mt-cyb, mt-nd5* and *mt-nd6* in relation to other loci, suggestive of LSMD hotspots. This is consistent with the previous report investigating LSMDs in non-tumorigenic brain samples that also showed 5’-breakpoints were most frequent in *mt-co1* and 3’-breakpoints were frequent in *mt-cyb* (with *mt-nd5* being the most affected, Hjelm et al. 2019). Large differences in sample number/types and sequencing and bioinformatics strategies likely contributed to the differences in absolute numbers of breaks, with (Hjelm et al. 2019) using short-read Illumina sequencing and their pipeline, “Splice-Break”. It is possible that sequence-specific features (e.g., repeat regions) and/or structural features (e.g., G4s) can predispose these (and other) regions of the mtDNA to deletion events. G4s are four-stranded nucleic acid structures formed by guanine-rich sequences in DNA and RNA. They are non-canonical structures, meaning they deviate from the typical double-helix structure of B-form DNA. G4s are formed by Hoogsteen hydrogen bonding of four guanine bases in a planar arrangement called a G-quartet, stabilised by a cation (e.g., potassium), which then stack on top of each other to form the four-stranded structure. They can exist in different topologies depending on the orientation of the guanines within the quartet, and can be found in various genomic locations, including telomeres, promoters, untranslated regions of mRNA, and mtDNA.

Consequently, we looked at the location of two LSMDs breakpoints relative to repeat regions and putative secondary structural motifs. These features have long been implicated in the formation of mtDNA deletions through slipped-strand mispairing, and replication stalling mechanisms (Phillips et al. 2017). Here, identical, or highly similar DNA sequences are mispaired during replication by the mtDNA polymerase, which slides along the template leading to the removal of the region between the repeats. Furthermore, replication may stall at non-canonical DNA structures such as G4s, causing it to become damaged through ds-breaks. This could be mediated by microhomology-mediated end joining (MMEJ), one of the main DSBR repair mechanisms in the mtDNA, which has been implicated in the formation of LSMDs. Since MMEJ requires micro-homologous regions of 5-22 bp in length, these imperfect repeat sequences may have determined the specific location where these LSMDs formed. Thus, a plausible mechanism for LSMD formation in cells/tissues involves genomic instability caused by G4 structures, replication fork slippage, transcription arrest, leading to the introduction of double-strand breaks (DSBs) via Endonuclease G that are then repaired via MMEJ, ultimately resulting in the formation of LSMDs (Dahal et al. 2022). This might be particularly true in the context of mtDNA, which is very G-rich and has fewer ROS-protective mechanisms and is therefore prone to destabilization and the formation of G4s (Aleksič, Podbevšek, and Plavec 2025).

The two LSMDs investigated more closely in this study were adjacent to or within either perfect or imperfect repeats, or potential G4-forming motifs, identified using QGRS mapper and AF3. In both cases scores were low, although AF3 suggests certain putative G4 sequences identified with QGRS Mapper are more likely to form in the presence of K+ ions. This is perhaps as expected, since metal cations coordinate the G-tetrads leading to the stabilisation of G4s (Phan 2010). While AF3 represents a significant advancement in macromolecule (particularly protein) structure prediction, its ability to predict structures is influenced by the availability of accurate experimental and training structures and/or evolutionary information, such as homologous sequences e.g., from the Protein Databank (PDB, (Berman et al. 2000)) and other databases. While there are some 70 G4-structures in the PDB, none are from mtDNA, which may have limited the ability of AF3 to predict structures with confidence for the LSMDs m.6130_15055del in the KSS samples and m.8469_13446del in fibroblasts_nd.

While it is possible that some G4-structures are more likely to form than others, based on specific sequences, the balance between ds-DNA and G4-stability and the chemical environment at a particular point in time, other “spare” G-tracts may too form into G4s, and may serve to mitigate the effects of the damaged G4s, preserving G4-function under stress conditions (Aleksič, Podbevšek, and Plavec 2025). The predicted G4-sequences in this study, yet unresolvable by AF3, might represent such “spare” regulatory elements. As more G4, especially mtDNA-G4, structures become available the ability of AF3 to predict structures accurately is set to increase and should shed light on the authenticity/importance of the other putative G4-sequences identified by QGRS Mapper in the future.

Nonetheless, these findings could suggest that the mtDNA deletions, including those detected by “NanoDel”, are not random, and that their formation is guided by both inherent sequence architecture and interactions with other factors. This includes ions that might be modulated within the mitochondria by the cellular microenvironment, thus contributing to our understanding of mtDNA instability in both health and disease. Interestingly, the two LSMDs investigated in more detail in our mitochondrial disease samples (this study) overlap with deletions observed in aging, metabolically active, and cancer tissues, as well as in neurodegenerative diseases, suggesting a shared vulnerability landscape shaped by mtDNA sequence motifs and structural constraints. These findings support the concept of “deletion hotspots” (Schon et al. 1989) as a unifying feature of mitochondrial dysfunction across multiple pathological states.

## Supporting information

NanoDel_SI

## Author Contributions

Conceptualization – REM (lead), Data Curation – REM, SCR and CFe (equal), Formal Analysis – REM, SCR and CFe (equal), Funding Acquisition – REM (lead), Investigation – CFe and REM (equal), Methodology – CFe, SCR and REM (equal), Project Administration – REM (lead), Resources – REM and SCR (equal), JP and CFr (supporting), Software – CFe (lead), SCR (supporting), Supervision – REM (lead), SCR and RB (supporting), Visualization – CFe (lead), REM (supporting), Writing – Original Draft Preparation – REM (lead), CFe (supporting), Writing – Review & Editing – REM (lead), CFe, CO, CG, JP, CFr, RB, SCR (supporting).

## Acknowledgments

We are thankful to Shelest, E. and Lundgren, A.P., who aided in the design of the script for clustering LSMD breakpoints in the “NanoDel” pipeline. We also gratefully acknowledge Beckett, A. and Dent, H., who helped with the library preparation and sequencing. CFe was supported by a University of Portsmouth Faculty-PhD studentship. For the purpose of open access, the author(s) has applied a Creative Commons Attribution (CC-BY) licence to any author accepted manuscript version arising from this submission.

