## Supplementary material for "NanoDel: Identification of large-scale mitochondrial DNA deletions using long-read sequencing": NanoDel_SI

### Supplementary Information:

#### *Supplementary materials and methods:*

*Cell cultures used as mtDNA amplification controls*

*Two human adult non-neoplastic astrocyte cell lines derived from the cerebral cortex (control) were obtained commercially (ScienCell research laboratories, #SC-1800) for use as mtDNA positive amplification controls. The 143B osteosarcoma and 143B rho-0 (a derivative of the 143B cells that have lost their mtDNA) cell lines were obtained commercially (Kerafast, ESA106), for use as mtDNA positive and mtDNA negative amplification controls, respectively (Table S2).*

*LR-PCR*

*Two-amplicon LR-PCR:*

*Whole mtDNAs (including any LSMDs) were amplified in two overlapping amplicons of 8,573bp and 9,128bp in length using human mtDNA specific primers in a 50µL reaction volume using the Expand Long Range dNTPack kit (Roche) (Lloyd et al. 2012)(Lloyd et al. 2015) (Keatley et al. 2019). The 50µL reaction contained 150nM (Amplicon 1) or 300 nM (Amplicon 2) of each primer, 1x PCR buffer, 500µM of each dNTP, 3% DMSO, 3.5U of polymerase and 8ng/µL of total DNA. The reaction was heated for 40 cycles with the lid heated to 103°C, 92°C (De Coster et al. 2018) initial denaturation, followed by 10 cycles of a 10 second 92°C denaturation, 15 second 58°C annealing and 10 minutes 68°C extension. This is followed by 30 cycles of 10 seconds at 92°C, 15 seconds at 58°C, 10 minutes at 68°C increasing by 10 seconds per cycle and a final extension at 68°C for 7 minutes. PCR products were assessed by Bioanalyser Fragment Length Analysis (1µL) using the Agilent DNA 12000 kit, and the Agilent 2100 Bioanalyser instrument, as described by the manufacturer's protocol. Products were then purified using the QIAquick® PCR purification Kit (Qiagen).*

*One-amplicon LR-PCR:*

*Each sample was also amplified as a single 16,266bp amplicon using the Takara LA Taq DNA polymerase kit (Takara Biosciences, RR002M). PCR reactions were completed in a 50μL reaction volume containing 200nM of each primer, 1x LA PCR buffer, 400nM each dNTP, 2.5U of LA Taq and 0.4ng/μL of total DNA input. The reaction was heated to 94°C for a 2-minute initial denaturation. It was then heated to 94°C for 10 seconds, 65°C for 15 minutes for 10 cycles followed by 94°C for 10 seconds and 65°C for 15 minutes increasing by 20 seconds per cycle for 30 cycles. Finally, they were heated to 72°C for 10 minutes for a final extension. Agarose gel electrophoresis using a 0.8% agarose gel was used to analyse approximate fragment lengths. PCR products were then purified using AMPure XP beads (Beckman Coulter, A63881) at a 2:1 (beads:PCR product) ratio and quantified using Qubit Fluorimetry (Thermofisher, Q33238).*

*Library Preparation and Sequencing*

*Library preparation was completed using the Native Barcoding Kit 96 V12 library kit (ONT, NBD112.96) according to manufacturer's specifications with small alterations. Individual amplicons underwent end preparation using the NEB Ultra End Prep and dA tailing module (New England Bioscience, E6053S). Individual amplicons were balanced in a 96-well plate to 8ng/ul with nuclease free water, and a total DNA quantity of 100ng. To each reaction 1.75μL Ultra II End Prep Reaction Buffer and 0.75μL Ultra II End Prep Enzyme Mix was added. The 96-well plate was sealed and incubated in a thermocycler at 20°C for 5 minutes followed by 60°C for 5 minutes.*

*End repaired samples were barcoded using the barcodes supplied in the NBD112.96 kit and the NEB Blunt T/A Ligase MasterMix (NEB, M0367S). Each barcoding reaction contained 5μL Blunt T/A MasterMix, 3μL nuclease free water, 1.25μL end prepped DNA and 0.75μL of each respective barcode. The barcoding reactions were then incubated again at room temperature for 20 minutes. Post-incubation 1μL of EDTA was added to each individual reaction to halt the reaction.*

*Samples were then multiplexed into a single 1.5mL tube and cleaned using AMPure XP beads (Beckman Coulter, A63880). A bead volume of 0.4x pooled sample volume was used for bead washing. The sample/bead mixture was incubated on a Hula mixer for 10 minutes at room temperature. The bead pellet was washed in 700μL of freshly prepared ethanol which was subsequently removed and discarded. After being allowed to dry, the bead pellet was resuspended in 35μL of nuclease free water to elute the DNA. This was incubated at 37°C for 10 minutes, being agitated every 2 minutes. The eluate was then retained.*

*Adapter sequences were ligated to the multiplexed DNA sample using the NEBNext Quick Ligation module (New England Bioscience, E6056S). This reaction contained 30μL of the DNA sample, 5μL of adapter mix II, 10μL of the quick ligation reaction buffer and 5μL of the T4 DNA ligase. Excess adapter mix was then removed with bead washing using a 0.4x reaction volume of AMPure XP beads. This was incubated on a hula mixer for 10 minutes at room temperature and then spun down and pelleted on a magnetic rack. The bead pellet was washed in 125μL of short fragment buffer (ONT, NBD112.96) twice, pelleted and then DNA was eluted in 15μL of elution buffer (ONT, NBD112.96), incubated at 37°C for 10 minutes agitated every 2 minutes. DNA was then quantified using a Qubit Fluorometer (Thermo Fisher Scientific, USA). Where possible the prepared library was then mixed with 37.5μL of Sequencing Buffer (SQB) and 25.5μL of Loading Beads (LB) (ONT, NBD112.96) to a concentration of 1.4ng/μL and a final volume of 75μL.*

*Artificial data generation:*

*Two sets of artificial FASTQ files were generated using VISOR (v1.1, (Bolognini et al. 2020). These were generated in two ways: 1) an ONT style with a longer average read length of ~8kb and a "Nanopore 2020" style error model, and 2) an Illumina style with 250bp length reads and an "Illumina" style error model. The first set (artificial data 1) contained controlled quantities of reads designed to emulate the 4977-bp deletion, often referred to as the "common deletion" or mtDNA4977, occurring between m.8470_13447, removing 5 tRNA genes and 7 genes encoding subunits of the MRC. These deletions were simulated in 75%, 50%, 25%, 5%, 1% and 0.25% of reads. The second set of artificial FASTQ files (artificial data 2) contained multiple deletions at nucleotides m.8471_13449del, m.6620_8440del, m.3541_5500del, and m.12538_15472del, all simulated in approximately 10% of reads. These files were used to evaluate and compare four separate alignment and deletion calling strategies highlighted below.*

##### *Alignment and LSMD analysis strategies:*

*In this study four pipelines were compared to analyse the efficacy of using a splice-aware RNA alignment (for DNA reads) vs. a traditional DNA alignment algorithm to identify deletions.*

*For "NanoDel" (Pipeline 1), compiled specifically for this study (outlined in Fig 1), reads were filtered further removing those with a read length <20bp and a Q-score of 10. Reads were aligned using minimap2 (v2.21, (Li 2018)) with the "splice" setting enabled but preferences for canonical splicing disabled. The BAM files were indexed using SAMtools (v1.7, (Li et al. 2009)). Analysis of junctions within reads was completed using the JWR_Checker python script from NanoSplicer (v1.0, (You, Clark, and Shim 2022)) and used to output a .csv file containing all junctions within reads. The .csv files were formatted using an R script and deletion breakpoints were clustered using sci-kit learn (v1.1.1, (Pedregosa et al. 2011)), DBScan (Ester et al. 1996) within a distance (eps) of 10, and a minimum cluster requirement of 2 samples. A* *mean value of each breakpoint in the cluster was then taken and reported back as the deletion, with a summation of the total reads for each deletion.*

*For "Splice-Break" (Pipeline 2), files were deduplicated and reformatted using BBmap (V38.18, BBTools (Bushnell 2020)). Reformatted paired end FASTQ files were then aligned using RNA aligner Mapsplice2 (v2.4, (Wang et al. 2010)) using the non-canonical double anchor setting. The "Splice-break".sh bash script (Hjelm et al. 2019) was then used for junction analysis, which created a .txt file containing deletion breakpoints.*

*Approaches for "Wood ONT" (Pipeline 3), and "Wood Illumina" (Pipelines 4) are based on those described in (Wood et al. 2019). "Wood ONT" (Pipeline 3) made use of Minimap2 (v2.21, (Li 2018)) with the DNA alignment "Map-ONT" setting. Structural variants in the resulting BAM file were then analysed using NanoSV (v1.2.4, (Cretu Stancu et al. 2017)), outputting a VCF file. Structural variants were then called from the VCF file using sniffles (v1.0.12, (Mahmoud et al. 2019)). "Wood Illumina" (Pipeline 4) aligned reads using BWA-mem (v0.7.17). Split and discordant reads were extracted from the .bam files using BWA-mem and SAMtools. Structural variant analysis was conducted using LumpyExpress (v0.2.14, (Layer et al. 2014), and the VCF file generated was genotyped using SVTyper (v0.7.1, (Chiang et al. 2015))). "Wood ONT" (Pipeline 3) typically uses the 'Canu' software to correct and trim the sequences prior to alignment with Minimap2. However, during tests on artificial FASTQ data, Canu removed all reads from the data set, due to aggressively down-sampling coverage. This read correction was likely confounded by the effects of heteroplasmy, leading to excessive reads being removed. Consequently, for the analyses of artificial data sets we also proceeded without using Canu.*

*The "NanoDel" pipeline is freely available for use on GitHub: https://github.com/uopbioinformatics/NanoDel.*

*Benchmarking and filtering:*

*For "NanoDel" (Pipeline 1), "Wood ONT" (Pipeline 3, (Wood et al. 2019)) and "Wood Illumina" (Pipeline 4,(Wood et al. 2019)), the number of deleted reads were normalised for each sample against a coverage benchmark to provide a relative read % for each LSMD. Coverage benchmark regions were established through the visible identification of a 500bp region with an even un-interrupted coverage across samples between positions m.500_1000del. "Splice-Break" (Pipeline 2, (Hjelm et al. 2019)) has an in-built benchmarking approach that uses two 250bp regions between positions m.957_1206del and m.15357_15606del to normalise the deleted reads and provide a relative read % for each LSMD between samples.*

*Additionally, for "NanoDel" (Pipeline 1) and "Splice-break" (Pipeline 2, (Hjelm et al. 2019)), a separate positional filter was included for deletion calls. Any deletions with a breakpoint occurring within 500 base positions of a primer boundary were removed. According to Hjelm et al. 2019, this position filter removes false-positive detection calls that occurred due to incorrect primer binding and putative deletions that occur within the spike-in coverage that can occur near the site of LR-PCR primers. For "NanoDel" (Pipeline 1), putative deletions needed to be supported by 2 reads or more to be reported or present in >0.1% of reads to be called. "Wood ONT" (Pipeline 3) and "Wood Illumina" (Pipeline 4) made use of the assembly tool Canu (v2.2, (Koren et al. 2017)(Jain et al. 2018)), specifically the correction and trimming functions. These functions replaced noisy read sequences with consensus sequences computed from overlapping reads, to decide which regions contain "high-quality" sequences, trimming the read to retain the largest "high-quality" chunk of sequence.*

*Repeat/ G4 sequence analysis and adjusted positions*

*Sequences flanking the 5'- and 3'-breakpoints of the LSMDs occurring at ≥50% of reads detected in the filtered "NanoDel" data (Pipeline 1) were analysed for repeat sequences and their potential to form G-quadruplexes (G4sG4s are implicated in diverse biological processes, including DNA replication, transcription, translation, and genome stability. SAMtools (Li et al. 2009) faidx was employed to extract 50bp for the repeat sequence analysis or 100 bp for the G4 analysis on either side of the respective 5'- and 3'-breakpoints from the rCRS (which is displayed as the L-strand), respectively.*

*For the repeat sequence analysis, the extracted sequences flanking the 5'- and 3'-breakpoints were aligned using ClustalW (Sievers et al. 2011)). Regions containing the longest stretches of contiguous perfect or imperfect repeated sequence were identified visually from the alignments and reported. For the G4 analysis, the reverse complements (representing the H-strand) of these extracted sequences were generated using the online Java Sequence Manipulation Suite (Stothard 2000)). The QGRS Mapper tool was used to predict the G4-forming sequences from both the original and reverse complement sequences, applying default settings of a maximum length of 30 bases and a minimum G-group size of 2 (Kikin, D'Antonio, and Bagga 2006)). QGRS Mapper provided either single or multiple G4-forming sequences along with a G-score for each sequence indicating its likelihood to form G4 structures. The G-score is based on the length and evenness of loop regions and the number of guanine tetrads, with higher (>30) scores suggesting a strong likelihood to form. A sequence of 30 bp could have a maximum score of 105, corresponding to the sequence 5'-GGGGGGTGGGGGGTGGGGGGTGGGGGG-3'. Where multiple putative G4-forming sequences were found, the one with the highest G-score was reported and chosen for investigation using AlphaFold 3 (AF3) in the presence and absence of a K+ cations. The formation of G4s is dependent on monovalent cations, including K+, which is often seen at high (~140mM) intracellular concentration (Sen and Gilbert 1988)). Where only a single G4-forming sequences was reported, this too was investigated using AF3.*

*AF3 is the most recent version of the Google DeepMind's artificial intelligence modelling system, and is able to generate 3D structures using nucleic acids sequences not just proteins (Abramson et al. 2024). Repeat regions are often used to annotate LSMDs. For example, mtDNA4977 has two flanking 13-bp direct repeats (8470-8482 and 13447_13459). However, whether the first or last base of the repeats is used for annotation can lead to slight inconsistencies in numbering, as can the numbering system used by the alignment algorithm. For the two LSMDs investigated in more detail (this study), we took the first base of both the 5'- and 3'-of their associated repeats and presented adjusted positions (Table S6).*

***Supplementary results:***

*NanoDel most accurately identifies the mtDNA4977 at low levels in simulated data*

*Relative read percentage (Poulton et al. 1993) of all reads in which the LSMD was found relative to benchmark region coverage (read%) was used here as a proxy for heteroplasmy. This metric was used to assess the sensitivity of each pipeline in its ability to detect LSMDs at different read percentage levels, as well as their accuracy when detecting the correct locations of 5'- and 3'-breakpoint pairs for each respective LSMD. Artificial data 1 were designed to reflect controlled ratios of reads containing mtDNA4977 (m.8471_13449del) and wild-type mtDNA, recapitulating the effects of heteroplasmy in silico. Visual inspection of coverage traces revealed that the expected LSMD was visibly identifiable at read % levels of ≥5% using all four pipelines (Fig 2 A-D).*

*"NanoDel" (Pipeline 1) and "Wood Illumina" (Pipeline 4) were the most sensitive to low heteroplasmy LSMDs, as they were the only two pipelines that identified m.8471_13449del within 20 base positions of its true 5'- and 3'-breakpoints at all simulated read% levels (as low as 0.25%). "NanoDel" (Pipeline 1) was able to identify 98.4% ± 22.9 (mean ± SD) of the simulated reads containing a LSMD at each respective deleted read% level (Fig 2 E), with an*

*average sensitivity of 0.926 ± 0.180, whilst maintaining a specificity of 0.992 ± 0.011 across all deleted read% levels. "Wood Illumina" (Pipeline 4) identified 138.5% ± 28.5 of deleted reads on average, over-representing the true level of the LSMD, with a higher average sensitivity of 0.995 ± 0.013, but also a lower specificity of 0.976 ± 0.043. In contrast, "Splice-break" (Pipeline 2) and "Wood ONT" (Pipeline 3) both failed to detect LSMDs at 0.25 read % levels (Fig 2 E).*

*The ability of each pipeline to accurately detect the true location of different LSMD breakpoints was also determined (Fig 2 F). "NanoDel" (Pipeline 1) was the most accurate at detecting 5'- and 3'-breakpoints in the least distance from the true simulated genomic position. "NanoDel" was able to identify each 5'- and 3'-breakpoint of m.8471_13449del within 1 ± 0 nucleotide position, representing the LSMD as m.8470_13450del at all deleted read% levels (Fig 2 F). This was due to "NanoDel" implementing 1-based rather than a 0-based indexing system. "Wood Illumina" (Pipeline 4) identified two pools of LSMDs with similar 5'-breakpoints but differing 3'-breakpoints, but breakpoint calls made by "Wood Illumina" (Pipeline 4) were 26.5 ± 35 away from their true genomic position (Fig 2 F). "Splice-Break" (Pipeline 2) was the second most accurate at detecting the true breakpoint locations within 1.3 ±0.19 positions, in deleted read% levels of ≥0.25% (Fig 2 F). Similarly, "Wood ONT" (Pipeline 3) was able to determine breakpoints within 4.2 ± 3.9 positions, in deleted percentage levels of ≥5% (Fig 2 F).*

##### *NanoDel most accurately identifies breakpoints and heteroplasmy levels in mixed deletion populations in simulated data*

*The second set of artificial data were designed to reflect a more complex mixture of four (including some overlapping) LSMDs (which were deleted between positions m.8471_13449del, m.6620_8440del, m.3451_5500del and m.12538_14572del), in 10% of reads, respectively (Fig 3 A-D). This was done to recapitulate more complex biological samples. Drops in coverage were visible within deleted loci following the alignment stages of each pipeline (Fig 3 A-C), except for "Wood Illumina" (Pipeline 4, Fig 3 D). The coverage generated by "Wood Illumina" (Pipeline 4) remained static, making it difficult to deduce the presence of LSMDs visually. However, using "NanoDel" (Pipeline 1), "Splice-Break" (Pipeline 2) and "Wood ONT" (Pipeline 3), stepwise drops in coverage were visible due to the presence of overlapping LSMDs. In general, these overlapping LSMDs obscure breakpoint boundaries making it difficult to discern their true location through visual inspection alone (Fig 3 A-D).*

*All pipelines, except for "Splice-Break" (Pipeline 2), could identify all 4 LSMDs within the mixed LSMD profile of artificial data 2. "Splice-Break" (Pipeline 2) was unable to identify the LSMD m.3451_5500del but identified the remaining three (Fig 3 F). "NanoDel" (Pipeline 1) and "Wood ONT" (Pipeline 3) were able to detect 97.2% ± 18.7 and 96.1% ± 19.2 of deleted reads, respectively (Fig 3 F), with no significant differences between the pipelines (P>0.99 and P>0.97 respectively, Fig 3 E). "NanoDel" (Pipeline 1) only displayed a false positive detection rate of 0.01 and a false negative detection rate of 0.08, while "Wood ONT" (Pipeline 3) displayed the same false positive rate of 0.01 and a similar negative detection rate of 0.09. Both Illumina centric pipelines (Pipelines 2 and 4), identified 36.7% ±24.5 and 144.2% ±3.6 of deleted reads on average, respectively (Fig 3 F) showing significant differences from the expected (p=0.0001 and p=0.001, respectively) (Fig 3 E). "Splice-Break" (Pipeline 2) displayed a much higher false negative rate of 0.63 while "Wood Illumina" (Pipeline 4) displayed a higher false positive rate of 0.06.*

*"Splice-Break" (Pipeline 2) was the most accurate at determining the correct locations of 5'- and 3'-breakpoints, identifying them within 1 ± 0 (mean distance from simulated breakpoint ± SD), however, it was only able to identify 3 out of 4 LSMDs (m.6620_8440del,*

m.8471_13449del, m.12538_15472del). "NanoDel" (Pipeline 1) was able to identify all LSMDs within artificial data 2, within 1.25 ± 1 nucleotide positions of their respective 5'- and 3'-breakpoints on average. In comparison, "Wood ONT" (Pipeline 3) and "Wood Illumina" (Pipeline 4) identified them within 1.38 ± 0.7 and 3.13 ± 1.5, respectively. "Wood Illumina" (Pipeline 4) also identified a small sub-population of 4 separate LSMDs with the same corresponding 5'-breakpoints as those simulated but differing 3'-breakpoints 82.2 ± 2.6 nucleotide positions away. This sub-population of LSMDs was not included in the previous average distance in which the pipeline could detect 5'- and 3' -breakpoints (Fig 3 G) as these were deemed to be different LSMDs resulting from false positive detection.

LSMDs revealed in clinically diagnosed mitochondrial disease samples following different LR-PCR approaches

Due to the sensitivity, consistency, and accuracy of "NanoDel" (Pipeline 1) in detecting LSMDs within the artificial data, "NanoDel" was then employed to detect LSMDs and determine their breakpoints in two small mitochondrial disease patient cohorts. MtDNA breakpoints were also determined in these samples using "Wood ONT" (Pipeline 3) for comparison, both with and without 'Canu'. The first cohort contained DNA from 3 tissue samples (muscle, kidney and cerebellum) from a KSS patient (Table S2B), and the second cohort contained DNA from 5 samples (cells_nd, muscle_RNASEH1muscle, muscle_TWNK, fibroblasts_nd, and muscle_TYMP) from different mitochondrial disease patients (Table S2B).

Following two-amplicon LR-PCR, full length wild type mtDNAs were visible as two-amplicons at the expected sizes of 8.5kb and 9kb in the muscle and cerebellum from the KSS patient, of cohort 1, but no smaller amplicons suggestive of deleted mtDNAs were visible (Fig S1A-B). Full length wild type mtDNAs were also visible as two-amplicons at the correct size in four out the five samples (cells_nd, muscle_RNASEH1, muscle_TWNK, fibroblasts_nd, but not *muscle_TYMP due to the lack of material being available) from the patients diagnosed with mitochondrial disease in cohort 2. However, unlike cohort 1, smaller amplicons suggestive of deleted mtDNAs were visible in two samples (muscle_RNASEH1 and cells_nd) but not fibroblasts_nd and muscle_TWNK using the first primer set, and only in three samples muscle_RNASEH1, fibroblasts_nd, and muscle_TWNK, but not cells_nd using the second primer set (Fig S1A-B).*

*Following one-amplicon LR-PCR of the KSS samples, full length mtDNAs were visible as one-amplicon at the expected size (16.3kb), as were shorter amplicons suggestive of deleted mtDNAs, in muscle and cerebellum (Fig S1C). Full length mtDNAs were also visible as one-amplicon in fibroblasts_nd, muscle_TWNK and muscle_TYMP along with multiple smaller amplicons suggestive of deleted mtDNAs. Muscle_RNASEH1 and cells_nd appeared to contain either multiple deleted mtDNAs or a single homoplasmic deleted mtDNA, respectively, as no full length mtDNAs were visible (Fig S1C).*

*NanoDel yields lower error rates, higher mean coverage and higher maximum read lengths in pipeline comparisons*

*The read metrics from the two and one-amplicon LR-PCRs were initially combined after alignment of the ONT reads with "NanoDel" (Pipeline 1) and "Wood ONT" (Pipeline 3), read quality and length filtering (Table S4A, B). We observed overall that the average mapping rate percentage (Fig 4A), using "Wood ONT" (Pipeline 3, with Canu) (99.2% ± 1.8, mean ± SD) was significantly higher than that which was observed using both "NanoDel" (Pipeline 1) (76.6 ± 30.2, p=0.001) and "Wood ONT" (Pipeline 3, without Canu) (75.0 ± 29.2, p=0.0001). No significant differences were identified between "NanoDel" (Pipeline 1) or "Wood ONT" (Pipeline 3, without Canu). However, this increased mapping rate appears to have come at the cost of average coverage as "Wood ONT" (Pipeline 3, with Canu) (Sievers et al. 2011).6 ±*

*31.5) showed significantly lower levels of coverage compared to "NanoDel" (Pipeline 1) (1662.7 ± 932.0,p=0.0001) and "Wood ONT" (Pipeline 3, without Canu) (1759.1 ± 982.1, p=0.0001), which is most likely due to Canu's correct and trim functions removing the vast majority of reads. Again, no significant differences were determined between "NanoDel" (Pipeline 1) and "Wood ONT" (Pipeline 3, without Canu) (Fig 4C). "NanoDel" (Pipeline 1) was found to have the lowest average error rate of the three approaches; however, it was not significantly lower than the others (Fig 4B). No significant differences in the average read lengths aligned were identified between each of the pipelines (Fig 4D). However, significant differences were determined between the maximum read lengths aligned. "Wood ONT" (Pipeline 3, with Canu) on average aligned reads with a shorter maximum length (10775 ± 3499) compared to "NanoDel" (Pipeline 1) (19276 ± 4592, p=0.0001), and "Wood ONT" (Pipeline 3, without Canu) (19276 ± 4592, p=0.0001) (Fig 4E).*

*NanoDel and one-amplicon LR-PCR has an increased mapping rate and lower error rate than Wood ONT*

*Comparisons between the sequencing and alignment quality metrics following one-amplicon and two-amplicon LR-PCR were conducted to assess the impact of these approaches on mapping rate, error rate, coverage, and read length. The proportion of reads mapped to the reference genome were significantly higher following one-amplicon LR-PCR using "NanoDel" (Pipeline 1) (85% ± 31.1%) compared with two-amplicon LR-PCR (65.7% ± 27.6, p=0.02, Fig 4F). Similarly, "Wood ONT" (Pipeline 3, without Canu) exhibited a significantly higher mapping rate following one-amplicon LR-PCR (82% ± 29.6), compared to two-amplicon LR-PCR (65.3 ± 27.6, p=0.04, Fig 4P). No significant differences were identified between one-amplicon LR-PCR and two-amplicon LR-PCR when using "Wood ONT" (Pipeline 3, with Canu) as 99.14% ± 2.1 and 99.14% ± 1.4 of reads were mapped respectively (Fig 4K).*

*The error rate using "NanoDel" (Pipeline 1) following one-amplicon LR-PCR (0.03 ±0.003) was significantly higher compared to two-amplicon LR-PCR (0.02 ± 0.002, p=0.02, Fig 4G). Similarly, the error rate using "Wood ONT" (Pipeline 3, without Canu) following one-amplicon LR-PCR (0.13 ± 0.13) was significantly higher compared to two-amplicon LR-PCR (0.02 ± 0.001, p=0.03, Fig 4Q), but not when using "Wood ONT" (Pipeline 3, with Canu) (Fig 4L). While the error rate using "NanoDel" (Pipeline 1) following one-amplicon LR-PCR was significantly higher than the two-amplicon LR-PCR approach, the magnitude of the increase was low (Fig 4G). Furthermore, "NanoDel" (Pipeline 1) error rate was still lower (Fig 4G) than the error rates seen using "Wood ONT" (Pipeline 3, with and without Canu (Fig 4L and 3Q). All pipelines also generated a higher average coverage following two-amplicon LR-PCR. However, the increases were not considered significant in comparison to the coverage generated following one-amplicon LR-PCR (Fig 4H, M, R). The increase in coverage following two-amplicon LR-PCR was expected due to the overlapping nature of the two-amplicons causing specific regions to have 2x more coverage. Unsurprisingly, both mean and maximum read lengths are higher following one-amplicon LR-PCR for each pipeline however in each incidence the increase is not considered significant (Fig 4 I, N, S, and J, O, T, respectively).*

*NanoDel and one-amplicon LR-PCR best validates known LSMDs in KSS patient tissues*

*In addition to the differences between the quality metrics (Fig 4, Table S4A-B), visible differences were also apparent upon inspection of the coverage traces generated post-alignment using reads from the one or two-amplicon LR-PCR of the KSS samples. Homogenous drops in coverage, indicative of a singular LSMDs, were clearly visible at ~m.6000_15000 following one-amplicon LR-PCR using both "NanoDel" (Pipeline 1, Fig 5A) and "Wood ONT" (Pipeline 3, without Canu, Fig 5B). This was unlike those amplified using the two-amplicon LR-PCR approach where increases in coverage could also be seen in the regions*

*~m.6500_7000del and ~m.1500_15500del, using both "NanoDel" (Pipeline 1) and "Wood ONT" (Pipeline 3, without Canu) (Fig 5C and D, respectively). These spikes were presumed to be the result of overlapping ends of PCR products causing inflations in coverage. As such, drops in coverage appeared to be between ~m.7000_15000del. In all instances no drops in coverage were evident using "Wood ONT" (Pipeline 3, with Canu), due to low overall coverage (Fig B and D). Using "NanoDel" (Pipeline 1), under stringent filters including removing deletions called within ±500bp of a primer boundary, and focussing on a read percentage level of 5% or more, just one deletion (m.8469_13446del) was identified in one KSS tissue (kidney) using two-amplicon LR-PCR (Table S5A). In the absence of filtering however, an additional 33, 16 and 6 deletions were identified in the muscle, kidney, and the cerebellum samples, respectively (Table S5A). Using the same stringent filters, one deletion (m.6130_15055del) was identified using "NanoDel" (Pipeline1) in all three tissues (muscle, kidney, and cerebellum) at differing read levels relative to the benchmark (112%, 57%, and 92%, respectively) of the KSS patient following one-amplicon LR-PCR (Table S5A). These were consistent with the coverage traces for these samples, and the breakpoints previously determined in these samples via Southern blotting (Poulton et al., 1993, Poulton et al., 1989, Poulton et al., 1995, Table S2B). On the other hand, the average read % of m.6130_15055del was higher in the muscle, kidney and cerebellum using "NanoDel" and not correlated with the heteroplasmy % previously determined using Southern blotting (47%, 54% and 23%, respectively). In the absence of stringent filtering, a further 2, 18, and 6 deletions were called using "NanoDel" in the muscle, kidney, and cerebellum samples of the KSS patient (Table S5A).*

*Using "Wood ONT" (Pipeline 3, with Canu), and a read % of 5% or above, a set of 'breakends' spanning positions m.1_16568del were identified in each sample (muscle, kidney, and cerebellum) (see Table S5B) following two-amplicon LR-PCR. It was suspected that these sets of 'breakends' signified the ends of the reads relative to the rCRS reference, as the amplicon boundaries did not align directly with the rCRS. This increased to two deletions per sample when Canu filtering was removed (m.6835_6932del and m.6969_7068del in muscle, m.6578_8878del and m.6740_8699del in kidney, and m.6612_6714del and m.6745_6888 in cerebellum) (Table S5B). Consistent with "NanoDel" (Pipeline 1), deletion (m.6130_15055del) was also identified using "Wood ONT" (Pipeline 3 with Canu) in all three tissues (muscle, kidney, and cerebellum) of the KSS patient following one-amplicon LR-PCR, in addition to a set of 'breakends' spanning positions m.1_16568del in kidney (Table S5B). Without Canu, m.6130_15055del was also apparent in the three tissues, and two additional deletions (m.216_16189del and m.315_16184del) were also apparent in the kidney sample of the KSS patient (Table S5B).*

*The read% determined for m.6130_15055del in muscle kidney and cerebellum were correlated between "NanoDel" (Pipeline 1) and "Wood ONT" (Pipeline 3 with (106, 24 and 65%) and without Canu (112, 57 and 93%).*

*The lack of amplicons following either one or two-amplicon LR-PCR (Fig S1), consistently poor mapping rate, average read length and coverage (Table S4) for the kidney sample across all pipelines, suggests poor quality. It is therefore important to consider the importance of the use of high-quality DNA when interpreting LSMDs through mtDNA sequencing methodologies. While the lack of full-length amplicons after one-amplicon LR-PCR (Fig S1) for the muscle_RNASEH1 and cells_nd samples could also be suggestive of poor quality, this is unlikely due to the amplification of full-length amplicons post two-amplicon PCR (Fig S1), and the presence of consistent mapping rates, average read lengths and coverage of these samples across pipelines (Table S4).*

*Fig 5 show the 5'- and 3'-prime gene involvement of the deletions for filtered and unfiltered data, respectively, for "NanoDel" and "Wood ONT" (Pipeline 3), and that two-amplicon LR-*

*PCR (Fig 5 I-L), but not one-amplicon LR (Fig 5 E-H), prevents deletion calling across the amplicon boundaries.*

*As the two-amplicon LR-PCR completely failed to detect the known deletion (m.6130_15056del) in the KSS samples (and cannot detect deletions across amplicon boundaries), subsequent analyses of the other mitochondrial disease samples focussed on “NanoDel” (Pipeline 1) and “Wood ONT” (Pipeline 3) using the one-amplicon LR-PCR.*

*NanoDel and one-amplicon LR-PCR yield new insights into LSMDs in mitochondrial disease patients*

*After alignment with “NanoDel” (Pipeline 1) and “Wood ONT” (Pipeline 3) without Canu, drops in coverage were visible following one-amplicon LR-PCR for all patients. Samples cells_nd and fibroblasts_nd showed stepwise drops in coverage characteristic of singular LSMDs. However, samples muscle_RNASEH1, muscle_TYMP and muscle_TWNK did not show distinct homogeneous drops in coverage necessitating other tools for identifying breakpoints (Fig 6A and C). The exception to this is after alignment with “Wood ONT” (Pipeline 3) with Canu, where no obvious drops in coverage were visible, due to the poor overall coverage (Fig 6B).*

*“NanoDel” (Pipeline 1) identified more LSMDs within this cohort compared to the number identified using “Wood ONT” (Pipeline 3), with and without Canu. However, only six LSMDs were found to occur after stringent filtering (m.800_9530del and m.496_567del in muscle_RNASEH1, m.8469_13446del in fibroblasts_nd, and m.5787_13921del, m.555_14749del, and m.3262_14412del in muscle_TYMP) in three of the mitochondrial disease patients muscle_RNASEH1, fibroblasts_nd and muscle_TYMP (Table S5A). When the stringent filters were removed a total of 534 putative LSMDs were identified across all five samples with an average 106.8±114.2 LSMDs per sample (Table S5A). Only 19 had a read %*

greater than 5%. Additional LSMDs identified with filtering removed (but a read level of 5% or greater) included, LSMDs in muscle_RNASEH1 (m.1666_16070del, m.2818_16068del, m.10184_16074del, m.457_15889, m.1766_16070del, m.1646_16033del,1664_15971del, m.3166_16072del, m.2059_16069del, m.2270_16070del), no additional LSMDs in fibroblasts_nd, additional LSMDs in muscle_TYMP (m.3270_16089del, and m.104_16071del), and a single LSMD in cells_nd (m.8636_16072del).

10 LSMDs were identified in three out of five samples (muscle_RNASEH1, fibroblasts_nd and muscle_TYMP), using "Wood ONT" (Pipeline 3, with Canu) occurring in 12.8% ± 4.6 of reads on average with 9 out of 10 LSMDs occurring in 5% or greater reads including m.806_9519del, m.1664_15972del, m.1672_16071del, and m.10186_16076del in muscle_RNASEH1, m.8469_13267del and m.8482_13447del in fibroblasts_nd, and m.562_14181del, m. 5789_13923del and m.6630_13994del in muscle_TYMP (Table S5B). This increased to 35 LSMDs, of which 17 occurred in 5% or greater of reads identified by "Wood ONT" (Pipeline 3, without Canu). This included additional LSMDs in muscle_RNASEH1 (m.496_568del, m.460_15890del, m.1668_16071del, m.1765_16071del, m.2058_16071del, m.2273_16071del, m.2819_16071del, and m.3166_16071del), no additional LSMDs in fibroblasts_nd, additional LSMDs in muscle_TYMP (m.105_16071del, m.3264_14414del, m.3272_16068del, and m.5788_13918del), and a single LSMD in cells_nd (m.8648_16073del) (Table S5B).

These deletion profiles are consistent with the "single deletion type" determined using traditional approaches for fibroblasts_nd and cells_nd and the "multiple deletion type" determined for muscle_RNASEH1 and muscle_TYMP previously (Table S2B, Fig S1).

Previously, other methods could not determine the breakpoints of the deletions in muscle_RNASEH1, although the major deletion observed by Southern blotting was sized at ~8.6kb (Table S2B). "NanoDel" (Pipeline 1) offers novel insights in relation to muscle_RNASEH1 that: (1) the major deletion is m.800_9530del (8.7kb), (2) a further deletion (m.496_567del) is present but has historically been missed, and (3) the deletions have a relative abundance (based on read%) of 19 and 11%, respectively. The mtDNA profile observed is consistent with impaired mtDNA replication in muscle_RNASEH1, with pathogenic variants in RNASEH1, encoding ribonuclease H1, causing autosomal recessive progressive external ophthalmoplegia (PEO) with mtDNA deletions (Reyes et al. 2015). RNASEH1 is an enzyme that degrades DNA-RNA hybrid structures such as those crucial for initiating mtDNA replication.

While multiple deletions were previously observed in muscle_TYMP, their breakpoints were not determined (Table S2B). "NanoDel" (Pipeline 1) suggests deletions at m.5787_13921del, m.555_14749del, and m.3262_14412del each with a relative abundance of 13%, 8% and 5%, respectively. The mtDNA profile observed is consistent with the diagnosis of thymidine phosphorylase deficiency in muscle_TYMP, leading to thymidine build up and damage to mtDNA, which is often linked with the co-occurrence of multiple deletions, depletion, and somatic mtDNA point substitution due impaired mtDNA replication. muscle_TYMP's diagnosis of thymidine phosphorylase deficiency, presenting as autosomal recessive mitochondrial neurogastrointestinal encephalomyopathy (MNGIE) (Marti et al. 2005), was previously confirmed by the detection of pathogenic variants in TYMP, encoding thymidine phosphorylase.

A single deletion m.8483__13459del was previously described in fibroblasts_nd (Table S2B). "NanoDel" (Pipeline 1) reported the exact 5'- and 3'-positions of the deleted segment as m.8469_13446del. These are both in fact mtDNA4977, as both have a deletion size of almost 4977bp, and involve regions around 8470_13459. In the former case (Table S2B) the last base of the 5'- and 3'-repeats have been used for annotation. "NanoDel" also suggests a relative abundance of 79% (relative to benchmark region coverage) which had not previously been determined.

*Similarly, a single deletion m.8649_16084del was previously detected at ~72-76% heteroplasmy by qPCR in cells_nd (Table S2B). The "NanoDel" (Pipeline 1) called very similar breakpoints m.8636_16072del (but only without stringent filtering), at a higher relative abundance of 100%. While a diagnosis was not noted for cells_nd, according to MitoBreak (the mtDNA breakpoints database (Damas et al. 2014)), m.8649_16084del is associated with ageing tissues (Wei 1992) (Volmering et al. 2016). While there is not an exact match for m.8636_16072del (7436bp), other LSMDs with breakpoints located at m.8636 have been found in epidermal keratocytes irradiated with UVB (Hwang et al. 2009) and m.16072 has 57 hits in MitoBreak, with clinical diagnoses including mitochondrial myopathy and PEO (Regan 1999)(Wanrooij et al. 2004).*

*A total of 207 LSMDs were identified in sample muscle_TWNK using "NanoDel" (Pipeline 1), all with a relative abundance <5%. The LSMDs with the highest read percentage levels: m.5788_13922del, m.5790_13922del and m.6327_13990del were found to have relative abundances of 4.2%, 3% and 2.3%, respectively. This was consistent with the range of LSMD sizes identified via Southern blotting and a multiple deletion type (Table S2B) and gel electrophoresis following one-amplicon LR-PCR (Fig S1), the impaired mtDNA replication often observed in patients like muscle_TWNK, who was previously diagnosed with autosomal dominant twinkle (TWNK) related Progressive External Ophthalmoplegia with mtDNA deletions (TWNK)(Spelbrink et al. 2001). TWNK is a mtDNA helicase, which unwinds ds-mtDNA necessary for mtDNA replication.*

##### *Mt-co1, mt-nd5, mt-nd6 and mt-cyb harbour breakpoints most frequently in mitochondrial disease samples*

*Within the mitochondrial disease samples "NanoDel" in combination with one amplicon PCR determined 5'-breakpoints occurred most frequently in in mt-co1 (n=18 in the KSS samples and n=100 in the other mitochondrial disease samples, Fig 5F and 5E, respectively). While 3'-breakpoints occurred most frequently in in mt-cyb and mt-nd6 in the KSS samples (n=9 and 3, respectively, Fig 5F) and mt-nd5 and mt-cyb in the other mitochondrial disease samples (n=150 and 140, respectively, Fig 6E).*

*LSMDs identified by NanoDel in mitochondrial disease tissues lie adjacent to repeat sequences and putative G4 structure-forming sequences*

*Common features found to be associated with the formation of LSMDs are repeat sequences (Hjelm et al. 2019) and non-canonical secondary structures in DNA formed by guanine-rich sequences, known as G4-quadruplexes (G4s, (Dong et al. 2014)).*

*Looking at the breakpoints identified for m.6130_15055del in the KSS samples andm.8469_13446del in fibroblast_nd, which occurred at ≥50% of reads in filtered "NanoDel" data, fell within 1-15bp of a repeat sequence identified within the rCRS (Table S6). Similarly, all these breakpoints fell within 2-48bp of a putative G4-structure forming sequence identified by the QGRS Mapper with G-scores ranging from 9 and 20 (Table S6). The exception here was m.8469_13446del , where the 5'-breakpoint and the repeat sequence fell inside a putative G4-structure forming sequence with a G-score of 20. All the putative G4-structure forming sequences were found on the H-strand (Tabel S6).*

*Although the G-scores of the putative G4-forming sequences were considered low (<30), indicating they were unlikely to generate G4-structures under physiological conditions (Kikin, D'Antonio, and Bagga 2006), we still sought to investigate their propensity to fold into G4 structures using AF3 (Abramson et al. 2024).*

*AF3 predicted multiple G4 structures from sequence candidates we identified with the QGRS Mapper. Consistent with the low G-scores, the AF3 scores were also generally low, possibly due to an absence of these specific topologies in the PDB (Berman et al. 2000), Table S6).*

*However, including ions appeared to increase the AF3 prediction scores as might be expected (Aoki and Murayama 2012) (Table S6).*

*LSMDs identified by NanoDel in mitochondrial disease tissues are associated with a range of healthy and pathological tissues*

*For the two deletions investigated in detail above, we provided adjusted "NanoDel" positions based on their associated repeats. We then used these ± 10bp for overlap analysis with the MitoBreak database. Interestingly, the deletions overlapped frequently with breakpoints in the MitoBreak database and were associated with both a range of tissues and clinical features, including (and in particular) aged, metabolically active (e.g. brain and muscle), and tumour tissues, and neurodegenerative diseases (Table S6).*

***Figures:***

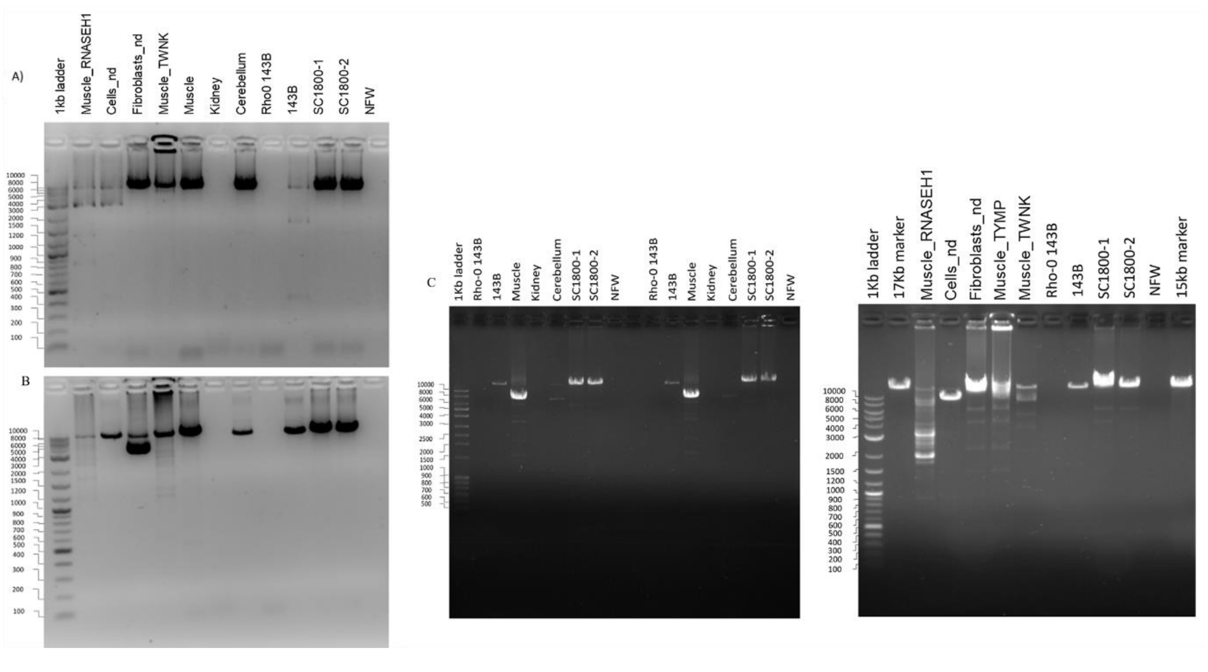


*Fig S1. LR-PCR amplification and agarose gel analysis of mitochondrial disease patient sample mtDNAs as two-amplicons overlapping primers A) CytbF and LongR/ Amplicon 1 (expected size ~8573bp), and B) LongF and Cytb/ Amplicon 2 (expected size ~9128bp); and as one-amplicon using almost back to back primers C) Alt Human LR-PCR F and Alt Human LR-*

*PCR R (expected size 16299bp). A 1kb DNA ladder and 17kb and 15kb DNA markers were used for sizing mtDNA bands. Muscle, kidney, and cerebellum samples were from a KSS patient, and cells_nd, Muscle_RNASEH1, muscle_TWNK, fibroblasts_nd, and muscle_TYMP were from patients with either confirmed or unconfirmed genetic diagnosis (Table S1B). SC1800-1 and 2 were from two human adult non-neoplastic astrocyte cell lines derived from the cerebral cortex, represent full length mtDNA + controls. 143B osteosarcoma and Rho-0 143B (a derivative of the 143B cells that have lost their mtDNA) cell lines, represent further full length mtDNA + control and a mtDNA – control, respectively. NFW (nuclease free water), represents no DNA input (PCR) – control. Smaller than full length mtDNAs are suggestive of LSMDs being present, and depending on the LR-PCR primers used, this reveals either "single" or "multiple" LSMD types.*

***Tables:***

*Table S1. S1 MtDNA variant types and clinical features*

*Table S2A. Cell culture details*

*Table S2B. Clinically diagnosed mitochondrial disease patient details*

*Table S3. Primer information*

*Table S4A. Pipeline 1 (NanoDel) read statistics for mitochondrial disease patients*

*Table S4B. Pipeline 3 (Wood Illumina) read statistics for mitochondrial disease patients*

*Table S5A. Pipeline 1 (NanoDel) LSMDs for mitochondrial disease patients*

*Table S5B. Pipeline 3 (Wood Illumina) LSMDs for mitochondrial disease patients*

*Table S6. Repeat regions and G4 sequences*

546
